## Supplementary figures and images for "Octopamine signaling from clock neurons plays dual roles in *Drosophila* long-term memory"

### Supplimental Figures

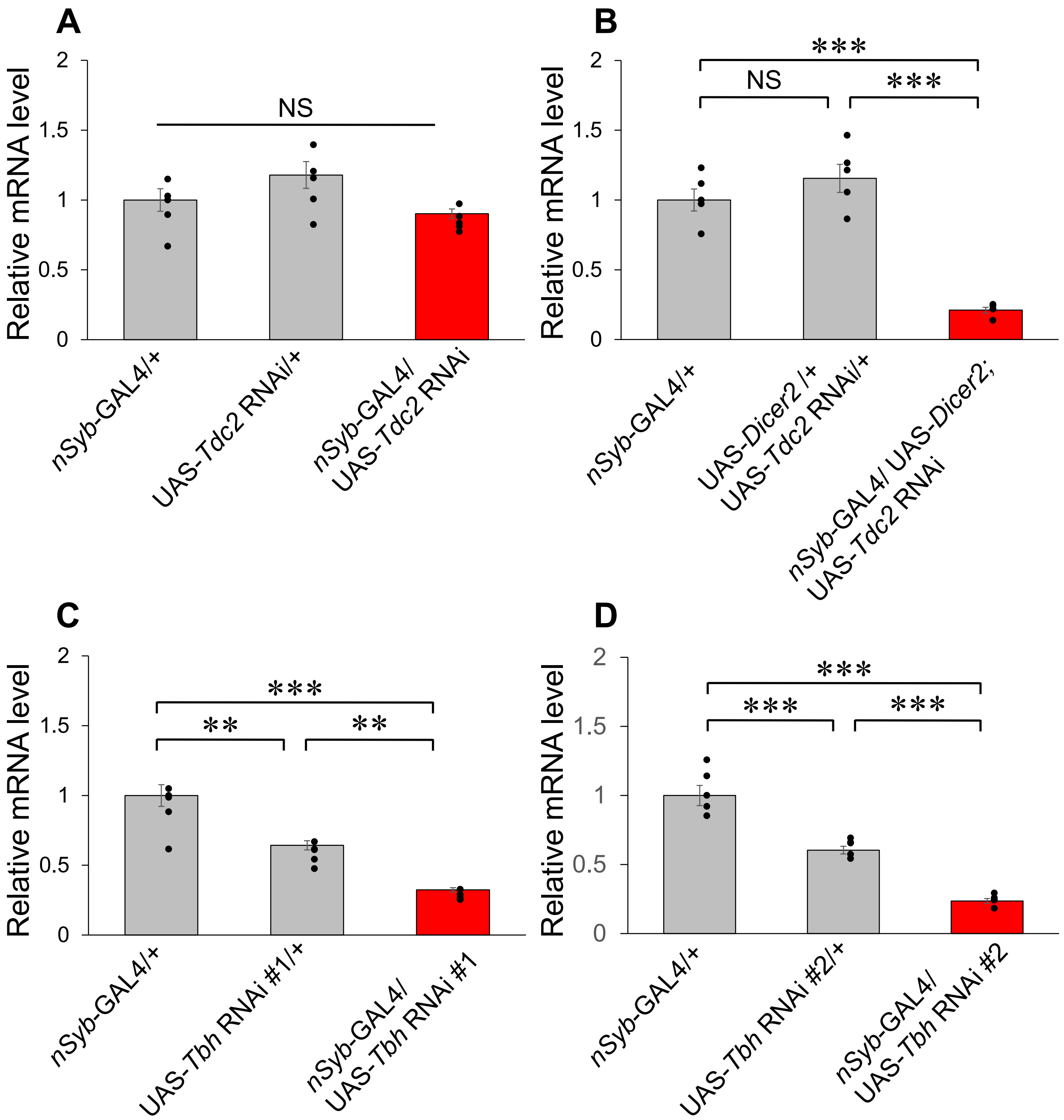

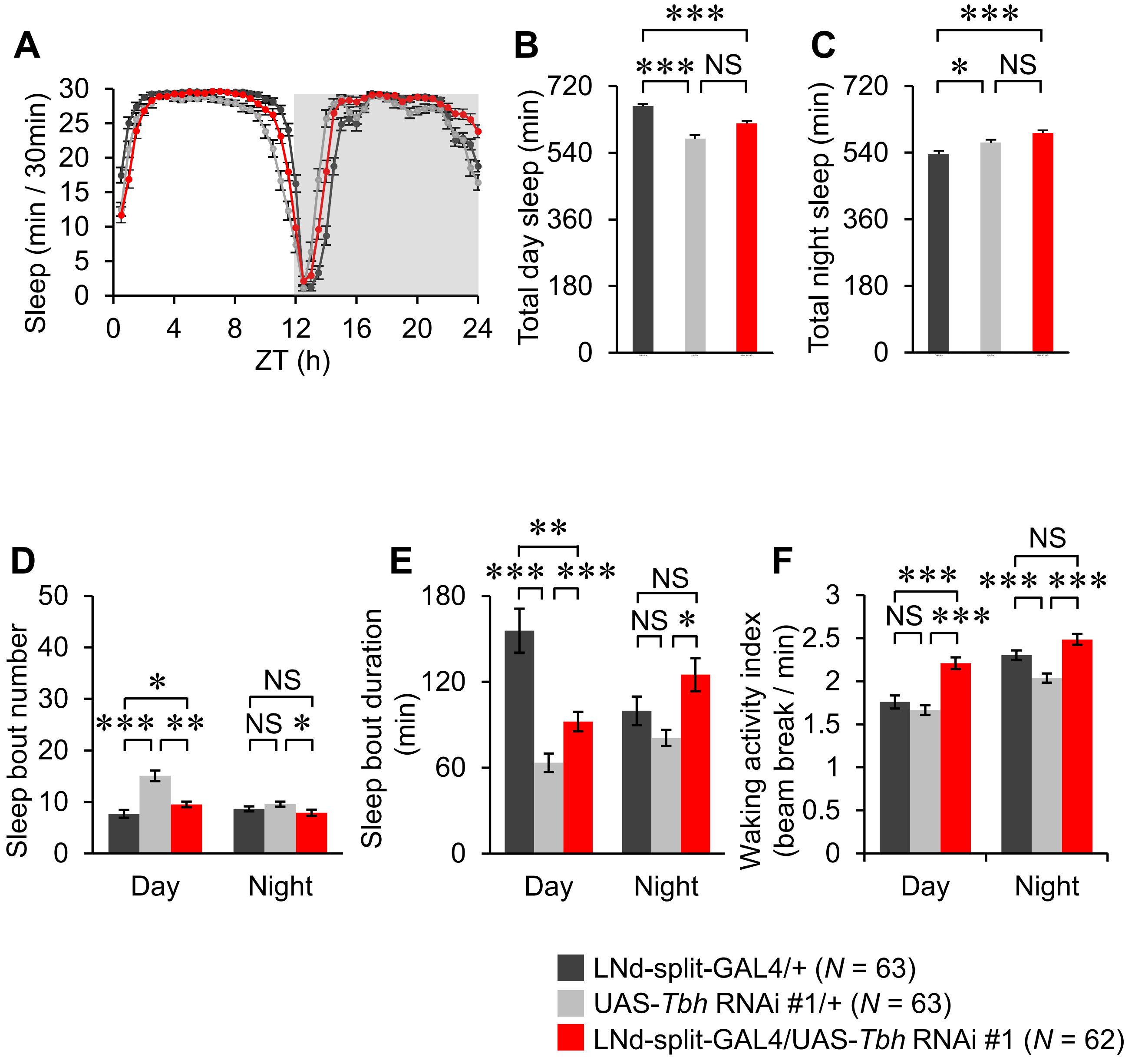

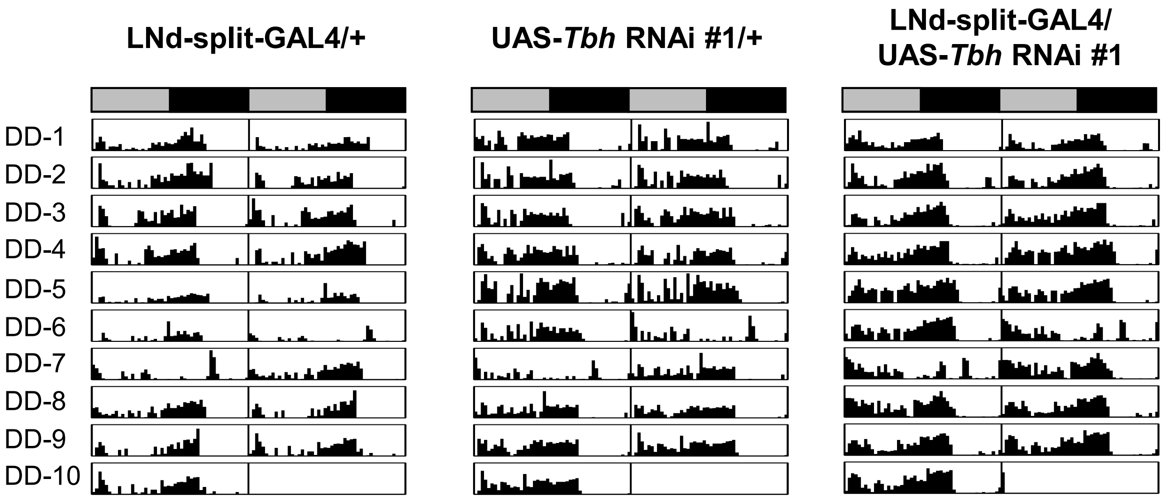


| | LNd-split-GAL4/+ | UAS-*Tbh* RNAi #1/+ | LNd-split-GAL4/ UAS-*Tbh* RNAi #1 |
|---|---|---|---|
| *N* | 63 | 63 | 62 |
| % Rhythmic | 74.6[NS] | 73.0[NS] | 79.0[NS] |
| Period (h) | 24.0 ± 0.06[A] | 24.1 ± 0.45[A] | 24.0 ± 0[A] |

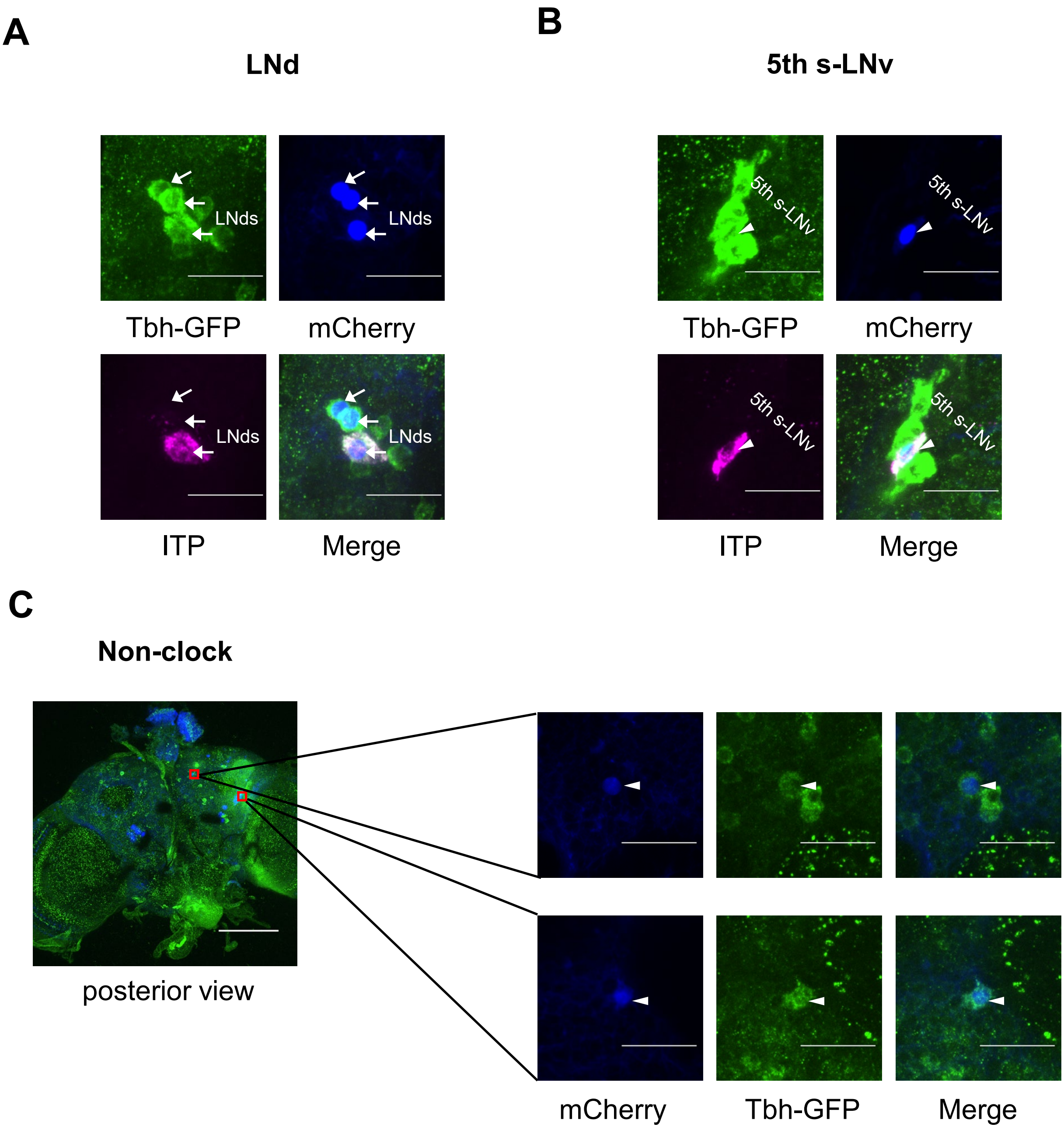

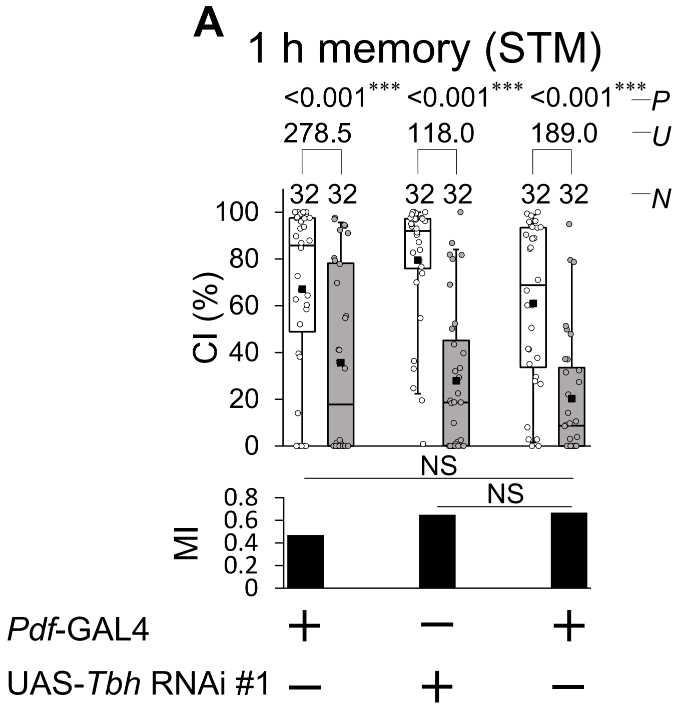

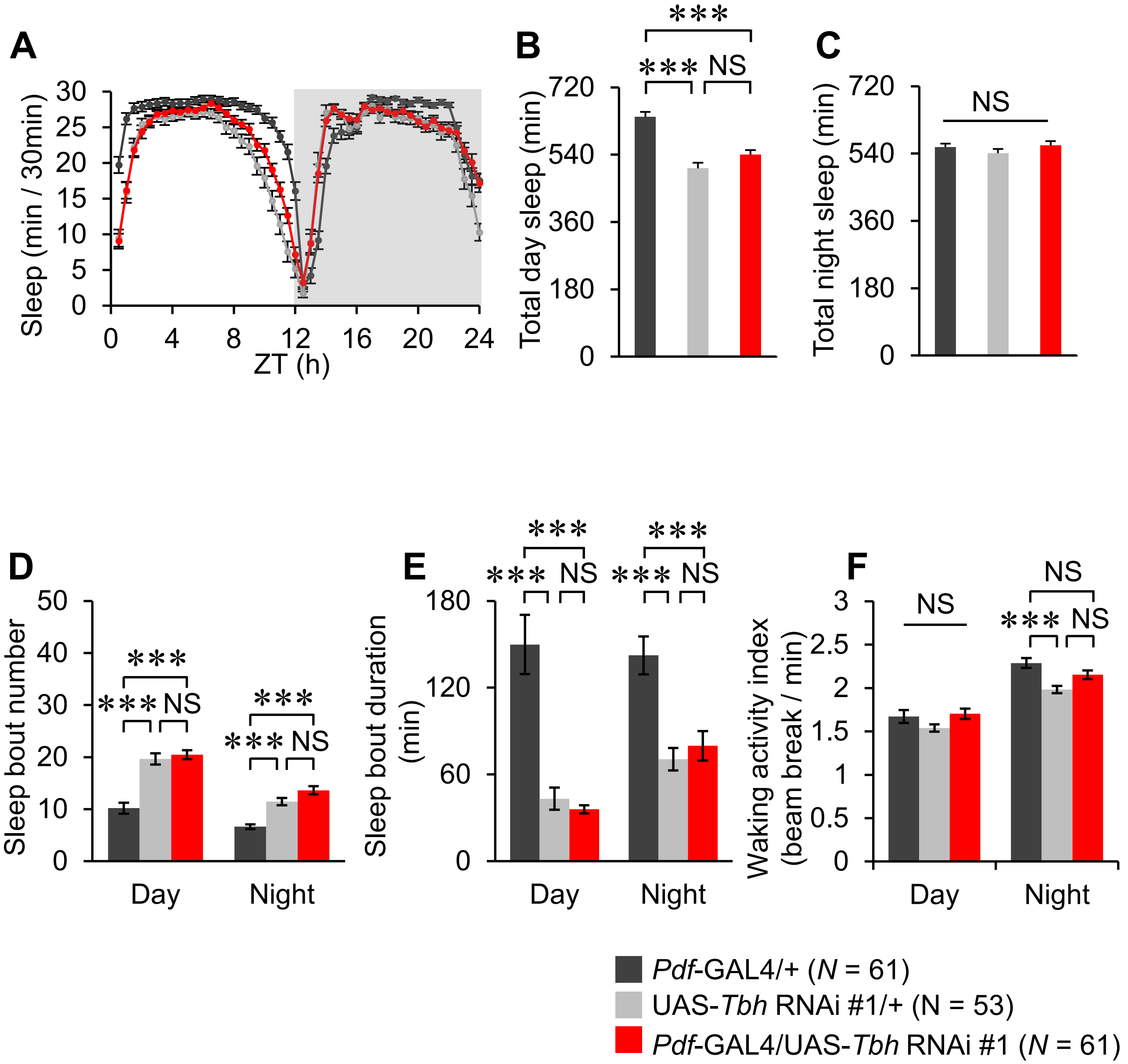
